# A Comprehensive Model of Blood Flow Restriction in the Post-surgical Rat

**DOI:** 10.1101/2024.03.27.586962

**Authors:** Andin Fosam, Susana Castelo Branco Ramos Nakandakari, Yuichi Ohashi, Hualong Bai, Julie O’Connell, Andrew Raines, Isabella Chavez Miranda, Alan Dardik, Christina Allen, Rachel J. Perry

## Abstract

Blood flow restriction (BFR) with low-load exercise (BFR-exercise) is an increasingly popular tool used to increase muscle strength and attenuate muscle atrophy, especially after injury or surgery. However, the mechanisms underlying BFR-mediated muscle growth are not well understood. Likely contributing to the mechanistic knowledge gap, rodent models of BFR-exercise have not been well described. In this methods paper, we demonstrate a comprehensive, clinically relevant protocol to establish BFR-exercise in awake rats. This protocol includes generating a muscle loss state via bilateral ACL-R, determining targeted blood flow occlusion pressures, and performing weighted hind-limb knee extension exercises with BFR. These methods can be used for further application in mechanistic and physiologic studies of BFR-exercise.

## INTRODUCTION

Blood flow restriction (BFR) is an increasingly popular tool used to increase muscle strength and attenuate muscle atrophy, especially after injury or surgery. BFR employs the use of an occlusion cuff to partially restrict arterial blood flow and completely restrict venous outflow to and from a distal muscle group^1–3^. When paired with light-weight exercise, BFR has demonstrated significant improvement in functional outcomes in humans experiencing muscle loss, including those recovering from orthopedic surgery^4–8^, aged populations^92,10–12^, and those with metabolic disorders^13–15^. Although the positive clinical outcomes are promising, a better understanding of the underlying physiologic mechanisms would allow for development of improved therapeutic protocols for treating muscle loss.

There is great potential for use of rodents to better understand the mechanisms underlying muscle hypertrophy following BFR-exercise. Indeed, rats are extensively used as a model system to investigate systemic and intramuscular adaptations to exercise^16^. Rat models allow for tight regulation of environmental and nutritional variables, access to functional analyses including *in vivo* tracers and pharmacologic agents, access to large and longitudinally collected tissue samples, and the ability for genetic manipulation. The use of rats to study resistance training has been generally explored through tail-weighted ladder climbing and incline treadmill walking^17–22^. There are several studies that have used rats to understand the effects of aerobic and resistance exercise on systemic and intramuscular metabolism^18,23–26^. Fewer studies have used rats to investigate the physiological effects and mechanism of action of BFR-exercise^2,27,28^. A compiled and comprehensive protocol to induce and utilize BFR-exercise *in vivo* in non-anesthetized rats has not yet been published.

From a clinical perspective, quadriceps muscle loss in the operative limb can reach ∼30-40% within 4 weeks following anterior cruciate ligament reconstruction (ACL-R) surgery in patients^29–33^. This atrophy is accompanied by significant functional deficits, including limited range of motion, and weakness, which results in prolonged recovery times^34–38^. BFR-exercise is used to attenuate muscle loss following common orthopedic surgeries, including ACL-R surgery in humans^4,6,39^. Rat models of ACL-R have been previously described to explore technical modifications and to trial engineered allografts of the ACL for ACL-R^40–44^. ACL-R as a model of post-surgical muscle loss for BFR-exercise in rats has not been extensively described but offers a clinically relevant foundation to study the hypertrophic effects of BR-exercise in rats. In this manuscript, we modify previously published rat models of ACL-R by using a commercially available engineered co-polymer allograft. Further, this paper describes completion of bilateral ACL-R, which is relevant for subsequent BFR-exercise procedures.

Achieving external BFR in rats has typically been done on anesthetized and carefully positioned rats^45^. Rubber bands and cuffs have been used to create transient arterial occlusion^22,46–48^, and direct surgical ligation is used for continuous arterial occlusion^49,50^. Indeed, external rubber bands are simpler to use than internal ligation, however it is difficult to achieve replicable occlusion pressures using this method. Fetal or digit cuffs (ranging from 1.6 to 1.9cm width) can generate more reproducible occlusions but have not yet been used to implement BFR in awake and exercising rats. This paper details a procedure using the 1.6cm digit cuff to induce targeted vascular occlusion in awake exercising rats. Further, it may serve as a resource for determining the therapeutic range of BFR occlusion pressures in rats.

Resistance training models in rats most commonly involve tail-weighted ladder climbing or incline treadmill walking^17–22,51^. These options employ similar muscle activation to traditional strength training modalities (*eg*. squats), but they are less applicable to studies of BFR due to their inability to generate hindlimb-specific movement, thus potentially introducing compensation by the non-targeted limbs. Other studies that combine BFR and resistance training in rats use low-current neuromuscular electrostimulation in anesthetized rats, which allows for careful control of occlusion pressure but only stimulates contraction of a single muscle^45,47,52^. Further, prolonged exposure to systemic anesthesia can cause reductions in body temperature and impact blood flow through distal vasculature, affecting accurate measurement of blood flow occlusion^53^. This does not fully reflect the contribution of synergistic muscles during the process of resistance exercise. Here, we describe our adaptation of a squat apparatus designed by Tamaki et al. that allowed for adjustable weighted knee extensions while isolating the hindlimbs during BFR exercise^54^.

Existing rat models of BFR exercise have provided key insight to the observed muscle hypertrophic adaptations. Establishing models that parallel clinical BFR-exercise procedures will allow for a much deeper understanding of its mechanism. The methods detailed in this paper lay a foundation for mechanistic studies of rehabilitative BFR-exercise in rats, including establishment of a ‘muscle loss state’, achievement of occlusion pressures within the BFR therapeutic range, and isolated knee extension exercise.

## METHODS

All animal surgeries were approved by the Yale Institutional Animal Care and Use committee (protocol #2022-20290). Male Sprague Dawley rats (6-10 weeks) were used for all animal experiments described below. The following step-by-step protocols detail each stage of the novel method we present here.

### Graft Preparation

1. Trim the 0.3mm wide FlexBand Plus co-polymer graft (Artelon®, Sandy Springs, GA) to a length of 3.0-3.5cm.
2. Using a scalpel or sharp fine scissors, trim the FlexBand graft lengthwise to a width of ∼1.0-1.5mm (about 1/3 of its width).
3. Attach a 5-0 Vicryl or PGA suture to each end of the trimmed graft using a Krackow stitch, leaving approximately 7-9cm suture tails on each end.

### Bilateral ACL Reconstruction Surgery

1. Inject subcutaneous pre-operative analgesia: buprenorphine XR (3.25 mg/kg) and bupivacaine (5 mg/kg)
2. Prepare the surgical area with sterile draping over a heated pad to prevent hypothermia.
3. Induce and maintain anesthesia by inhalation of 2% isoflurane with 2 L/min oxygen using the VetFlo Anesthesia Apparatus (Kent Scientific Corporation, Torrington, CT) for the duration of the surgery via induction chamber and nose cone respectively. Adequate anesthesia is confirmed by lack of response after applying sharp pressure to hindfoot with fingertip.
4. Remove hindlimb fur from both hind limbs (from hip joint to ankle) using depilatory cream (Nair) and then clean with 70% ethanol.
5. Place sterile draping over the animal leaving only the operative limb exposed.
6. With the knee in extension, make a 2-cm long vertical incision with a scalpel medial to the patella, centered at the level of the patella (Fig 1A).
7. Retract skin the laterally until the incision is centered over the knee joint. Slightly flex the knee and use the scalpel to open the parapatellar joint capsule by cutting directly medial to the patella and extending proximally to the level of the musculotendinous junction of the quadriceps and distally to the level of the patellar tendon insertion on the tibial tubercle. Note: Care should be taken to avoid transection of the medial collateral ligament (more medial), the patellar tendon (more lateral), or quadriceps tendon (more lateral).
8. With the knee flexed, use a pair of fine forceps to release and translate the patella laterally, exposing the inside of the knee joint. This should afford clear visualization of the intercondylar notch and femoral condyles (Fig 1B).
9. Transect the ACL in the intercondylar notch using a scalpel or pair of fine scissors. Successful transection of the ACL is confirmed by positive anterior drawer test by translating the tibia forward on the distal femur.
10. Using a power drill, place a 1.4-1.8mm stainless steel k-wire on the ACL origin in the intercondylar notch and drilled superolaterally through the lateral femoral condyle and exit on the lateral aspect of the femur. Ensure a patent and adequately dilated bone tunnel by carefully drilling back and forth within the bone. This will create a clear passage for the graft.
11. Thread a 1.4mm Keith needle (or tie a 3-0 or smaller Vicryl suture) with the one end of the suture tails of the prepared graft. Pass the Keith needle (or 3-0 Vicryl suture) through the femoral bone tunnel, pulling one end of the graft through the bone.
12. Secure the tails of the graft out of the way using a needle driver or mosquito snap (or similar tool) prior to creating the tibial bone tunnel.
13. Place a 1.4-1.8mm stainless steel k-wire on the ACL insertion point on the tibial plateau and drill anteromedially to the anteromedial proximal tibia.
14. Thread a Keith needle (or tie a 3-0 Vicryl suture) with the distal suture tails of the graft. Pull the needle with the attached graft through the tibial bone tunnel. Note: secure the proximal suture tails of the graft on the femoral side with a snap to avoid displacing it as the distal end is being pulled through the tibial tunnel.
15. With the knee in full extension, manually tension the graft and using a 4-0 Vicryl suture to fix the femoral end of the graft to the surrounding periosteum with a figure-of-eight stitch. Similarly, secure the tibial end of the graft to the surrounding periosteum or soft tissue with a figure-of-eight stitch. Remove excess graft and suture with scissors, leaving 0.5-1 mm on each end past the securing stitch (Fig 1C).
16. Flush the joint capsule with saline. With the knee in full extension, reduce the patella and close the medial joint capsule securely with 4-0 Vicryl or PGA suture.
17. Close the skin with 5-0 Vicryl or PGA suture and seal the wound with wound glue (Vetbond, 3M, Japan) (Fig 1D).
18. Remove the surgical draping from the operative hindlimb an inject 500uL saline subcutaneously at the level of the abdomen.
19. Place clean draping over the animal, exposing only the contralateral hindlimb.
20. Repeat steps 5-17 on the contralateral hindlimb.
21. Inject 500uL of analgesic (carprofen, 5mg/kg) into the intraperitoneal (IP) space. Stop the flow of anesthesia and remove the nose cone from the rat. Place the rat in a clean cage under fluorescent lighting. 5 mg/kg carprofen is also injected on post-operative days 1 and 2 for pain control.

**Figure 1.**
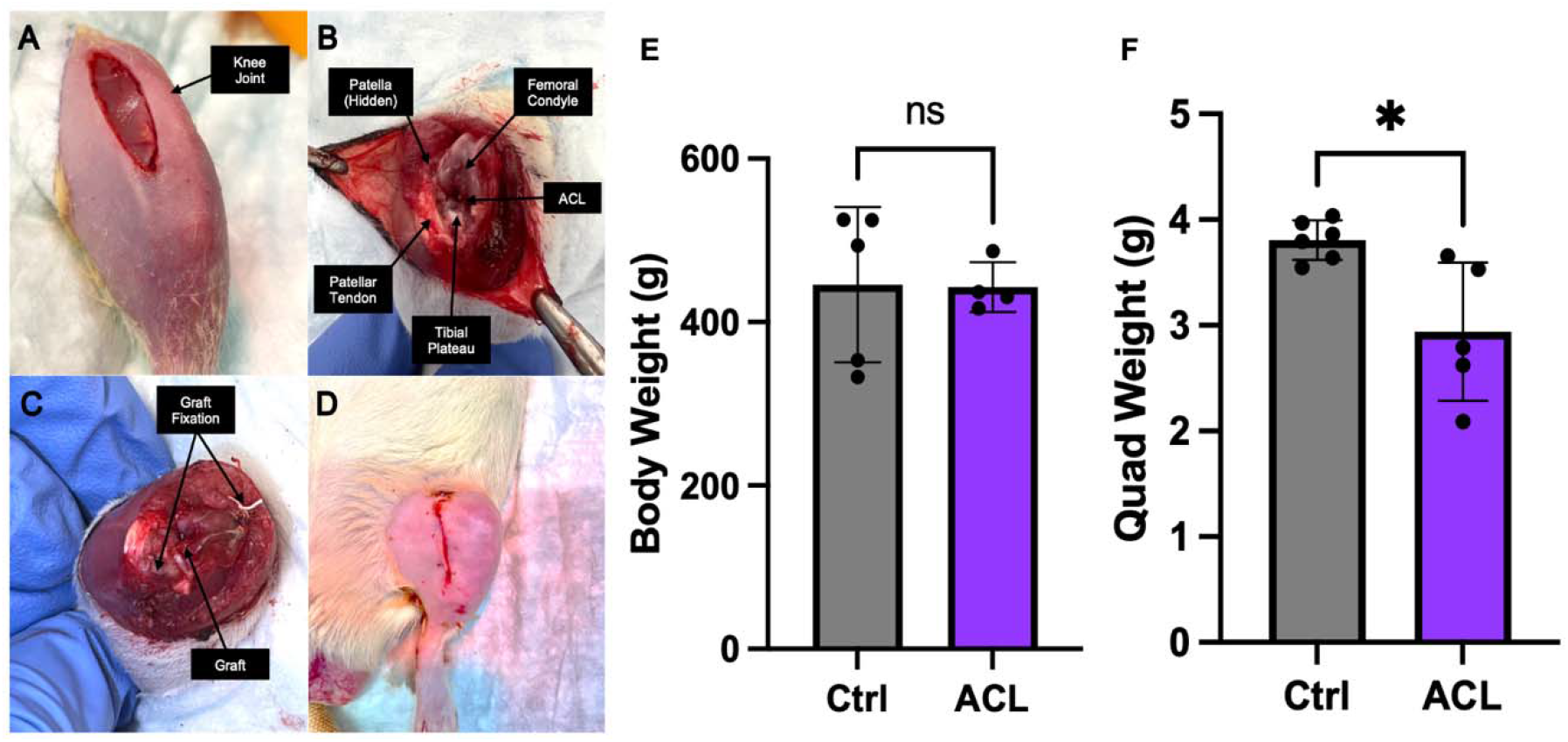
Muscle loss following bilateral ACL-R in adult rats. (A) Skin incision on the hindlimb medial to the patella. (B) Visualization of knee joint capsule with the patella translated laterally to view joint structures. (C) Fixation of engineered allograft of the ACL. (D) Closure of the skin incis on. (E) Bodyweight and (F) averaged quadriceps weight in non-surgical control rats as compared with ACL-R rats following a two-week recovery. *P<0.05. Male rats were used to generate data in this figure.

Throughout this procedure, it is necessary to monitor the respiratory rate of the rat throughout the course of the surgery. If breathing becomes shallow, the isoflurane flow rate may need to be adjusted. Additionally, the size of the K-wire can vary depending on size of the rat. We found that the 1.4mm K-wire is appropriate for rats ∼300g and the 1.6mm or 1.8mm k-wire for rats >400g.

### Cuff Placement Protocol (day of BFR-exercise intervention)

1. One day prior to occlusion cuff placement, anesthetize the rat and remove all fur from the hindlimb (or bilaterally, depending on the experimental objectives) using depilatory cream.
  a. Be sure to rinse the skin with 70% ethanol or normal saline to reduce skin irritation post-hair removal.
2. Line the underside of the cuff (the side that will be in contact with the skin) with double sided tape.
3. Place the rat in the harness and secure the harnessed rat to the squat apparatus.
4. Ensure that the cuff is fully deflated and attached to the sphygmomanometer.
5. While gently restraining the rat by hand, place the cuff around the proximal hindlimb, ensuring that it is secured above the knee joint.
6. Release the rat from the gentle hand restraint and watch for slipping or movement of the cuff, adjust as necessary.

### BFR Exercise Protocol

1. 7-10 days prior to the start of the BFR exercise protocol, rats undergo the following acclimation procedure: 1-2 days to freely roam with a clear box (with space along the perimeter for oxygen flow) containing the squat apparatus for 30 minutes to 1 hour; 1-2 days to roam the space containing the squat apparatus while wearing the harness for 30 minutes to 1 hour; 1-2 days strapped to the squat apparatus with no loaded weight for 7-10 minutes; 1-2 days strapped to the squat apparatus with loaded weight up to 15% bodyweight; 1-2 days strapped to the squat apparatus with loaded weight of 30% bodyweight. Note: Acclimation process can be incentivized with peanut butter.
2. Prior to the start of the BFR exercise protocol:
  a. Prepare the squat apparatus by loading the weights to the vertical pin until the desired load is reached. The squat apparatus used includes an 80g arm.
  b. Acquire any baseline blood or tissue samples prior to starting exercise.
3. Place the harness on the rat and tighten the straps. Be careful to not disrupt or occlude any catheters, if applicable.
4. Attach the harnessed rat to the squat apparatus. Ensure a secure attachment. Adjust the “rest position” height to achieve full knee flexion. The front limbs of the rat should not be allowed to contact the ground. If they do, raise the starting position.
5. Attach the occlusion cuffs to the rat as detailed above in the ‘Cuff Placement Protocol’ section.
6. Inflate the occlusion cuffs to a pressure within the therapeutic threshold (40-80% of the total limb occlusion pressure, ideally as determined by ultrasound) using a handheld sphygmomanometer (see Blood Flow Occlusion Protocol below).
7. Observe movement of the rat from the initial rest position to full knee extension.
  a. Knee extension can be prompted with an acute stimulus (ex: tail shock, compressed air targeted at the rectum).
  b. This model allows for variability in exercise regimens (eg. 3 sets of 10 repetitions, 30 repetitions in 10 minutes, etc)
8. After completion of the desired repetitions, unfasten the rat from the squat apparatus, remove the cuff and the harness, and return the animal to the cage.
9. Proceed with data collection as per the experimental design.

### Muscle Biopsy Protocol

#### Fine Needle Muscle Biopsy Protocol

1. The day prior to the start of the experiments, fully remove the fur of both hindlimbs from hip joint to ankle with depilatory cream. Once the fur is removed, cleanse the skin with 70% ethanol.
2. During the day of the procedure the hindlimb of the rat will be cleansed with 70% ethanol. Once dry, the rat will be secured in a supine position by gentle handheld restraint.
3. Anesthetize the skin of the hindlimb from the hip to knee joints using topical lidocaine 5% cream.
4. Place the quadriceps in a lengthened position by extending the hip and flexing the knee.
5. Orient and advance a 22-gauge spring-loaded core biopsy needle (Tenmo, Merit Medical Systems, South Jordan, Utah) into the anesthetized hindlimb at the level of the medial quadriceps.
6. Once positioned, release the spring-load function, closing the core trap area and capturing a small amount of muscle tissue.
7. Retract the needle from the skin.
8. Repeat steps 2-7 as needed for the specific experimental design.

### Blood Flow Occlusion Protocol

The Vevo 3100 system (Visualsonics Inc., Toronto, ON, Canada) paired with the UHF46x transducer probe was used in all experiments to measure changes in blood flow with varying degrees of external occlusion (0, 40, 60, 80, 100, 120 mmHg). Rats were anesthetized with isoflurane and positioned supine on the ultrasound table. Fur was removed from the hindlimb and abdomen with depilatory cream. The hindlimb was held in extension while a 1.6cm digit cuff (Hokason Inc., Bellevue, WA) was secured on the hindlimb above the knee joint. The external iliac artery was identified by first locating and following the abdominal aorta inferiorly until reaching the bifurcation point of the right and left common iliac artery. The right (or left depending on the experimental design) common iliac artery was followed until its bifurcation into the internal and external iliac artery (Fig 2A). Measurements of the external iliac artery were acquired 1-2 cm below the bifurcation point (or as distal as possible before the probe encountered the external cuff). In the short axis view (probe perpendicular to vessel), the diameter (mm) of the external iliac artery was measured at this location. Velocities (mm/s) were obtained in a long axis view (probe parallel to vessel) at the same anatomical position. Relative occlusion was calculated as the ratio of the vessel diameter at a given occlusion pressure to the vessel diameter at baseline (0mmHg occlusion pressure).

**Figure 2.**
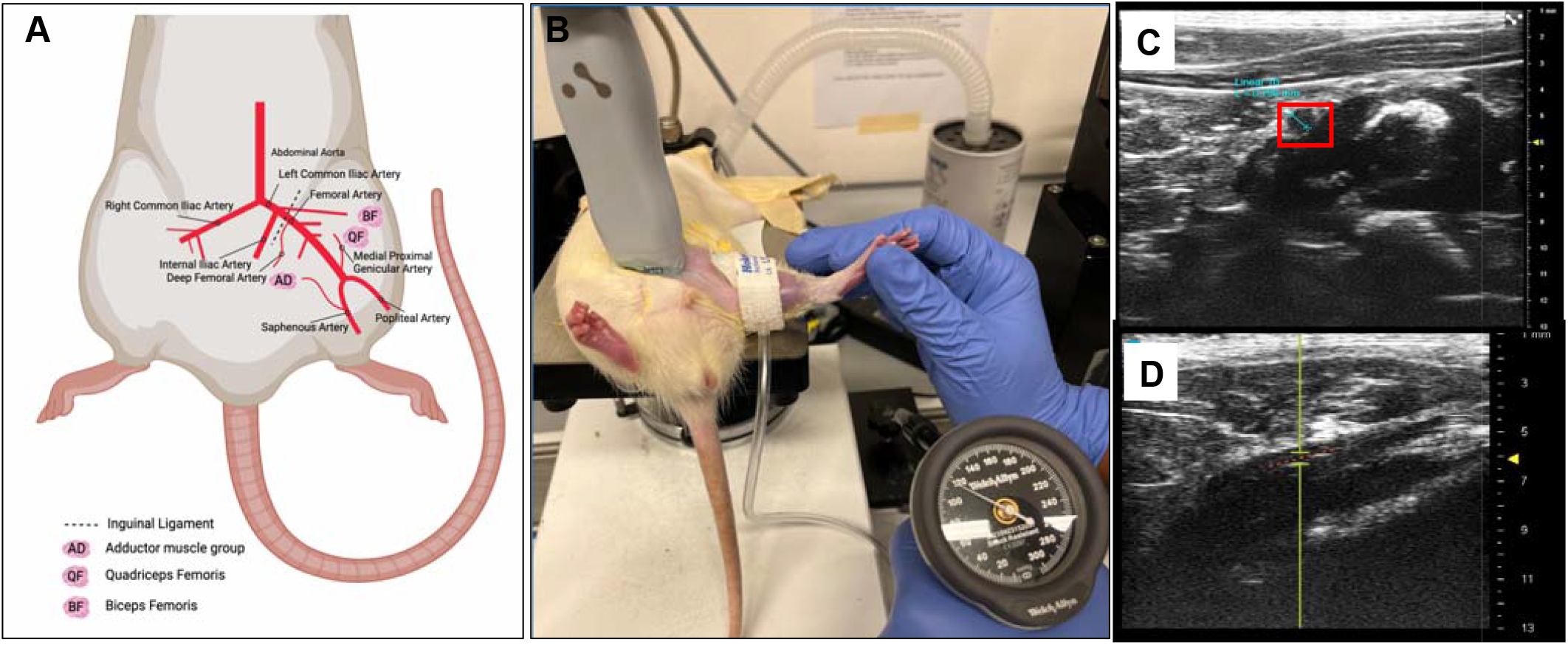
Generation of blood flow restriction in rats. (A) Rat hindlimb vasculature schematic depicting placement of ultrasound probe adapted from Aref, Zeen et al. Intl J Molec Sci. 2019. Created in BioRender. (B) Photo representation of occlusion cuff and probe placement, as well as rodent positioning to measure blood flow through the external iliac artery on an anesthetized rat. Cuff pressure set to 120 mmHg. (C) Doppler ultrasound image of the external iliac artery (red box) in short axis view and (D) long axis view (red dotted line) using the UHF46x probe.

### Statistics

Data are presented as mean ± standard deviation unless otherwise specified. Differences between two groups are calculated by the 2-tailed unpaired Student’s t-test (when comparing two groups).

## RESULTS

### ACL Reconstruction Surgery as a model of post-surgical muscle atrophy in rats

A total of 20 adult male Sprague Dawley (SD) rats (aged 6-10 weeks) underwent bilateral ACL reconstruction surgery with a mean operative time from incision to closure of the surgical wound of 40 minutes. There were no intra-or post-operative complications. Immediately following and one to two days post-surgery, the rats were observed to partially weight-bear on the surgical hindlimbs but were actively moving around the cage. By one-week post-surgery, the rats were fully weight-bearing on their operative hindlimbs with no appreciative limp. No wound abnormalities were observed, including bleeding, draining, erythema, swelling or dehiscence. The rats did not demonstrate any abnormalities in feeding, urinating, defecating, or sleeping in the post-operative period.

Fifteen of the 20 rats as mentioned above were used solely for the purpose of protocol development for this manuscript. These rats were euthanized approximately 14 days following surgery. The exercised rats were not included in the following analyses. At the time of euthanasia, bodyweights of the control and ACL-R rats were similar (Fig 1E). Both quadricep muscles were harvested and weighed immediately after euthanasia. Quadricep weight, normalized to bodyweight, was lower in the ACL-R rats compared to the non-surgical control group (Fig 1F).

### Rat model of BFR occlusion pressures

Digit cuffs (1.6cm) were placed on the hindlimb of the rat above the knee joint after complete removal of the fur using depilatory cream. Cuffs were secured to the skin using double-sided adhesive tape for the exercise. The external iliac artery was probed on the proximal hindlimb, above the occlusion cuff (Fig 2B). The vessel was visualized in both the short and long axis views (Fig 2C, 2D). In the short axis view, external iliac artery diameter is measured at 0, 40, 60, 80, 100, and 120 mmHg (Fig 3) in a prototypical rat. A vessel diameter of 0.867mm was observed at 0 mmHg. The smallest vessel diameter of 0.663 mm was observed at 80 mmHg, approximately 24% vessel occlusion.

**Figure 3.**
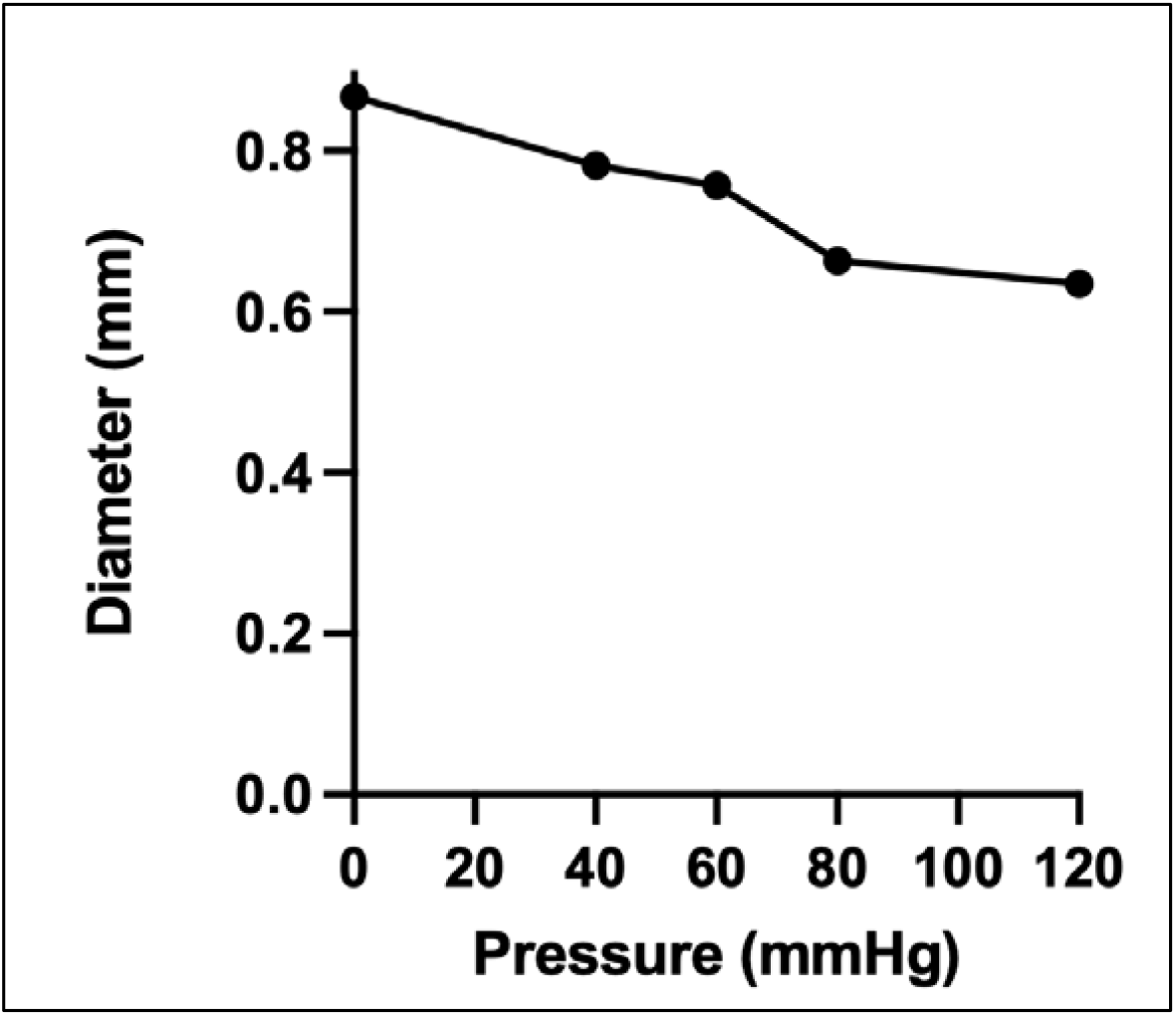
Ultrasound analysis of external iliac artery. Diameter in mm is measured at 0, 40, 60, 80, and 120 mmHg over the external iliac artery on the right hindlimb of a single rat.

### BFR exercise protocol

A total of 10 rats (aged 6-10 weeks) were used to develop the BFR-exercise method. The acclimation period on average lasted 7 total days. Fig 4A demonstrates approximately day 4 of the acclimatation during which the rat freely explores the squat apparatus while wearing the harness. Rats achieve full knee flexion at rest (Fig 4B) and full knee extension (Fig. 4C) to indicate one full repetition. Loaded resistance up to 30% of the rat bodyweight was achieved by adding weighted washers to the 80g arm of the squat apparatus. The rats had an average bodyweight of 421.6 ± 38 grams and squatted against an average of 122 ± 13 grams to achieve a resistance of 30% bodyweight.

**Figure 4.**
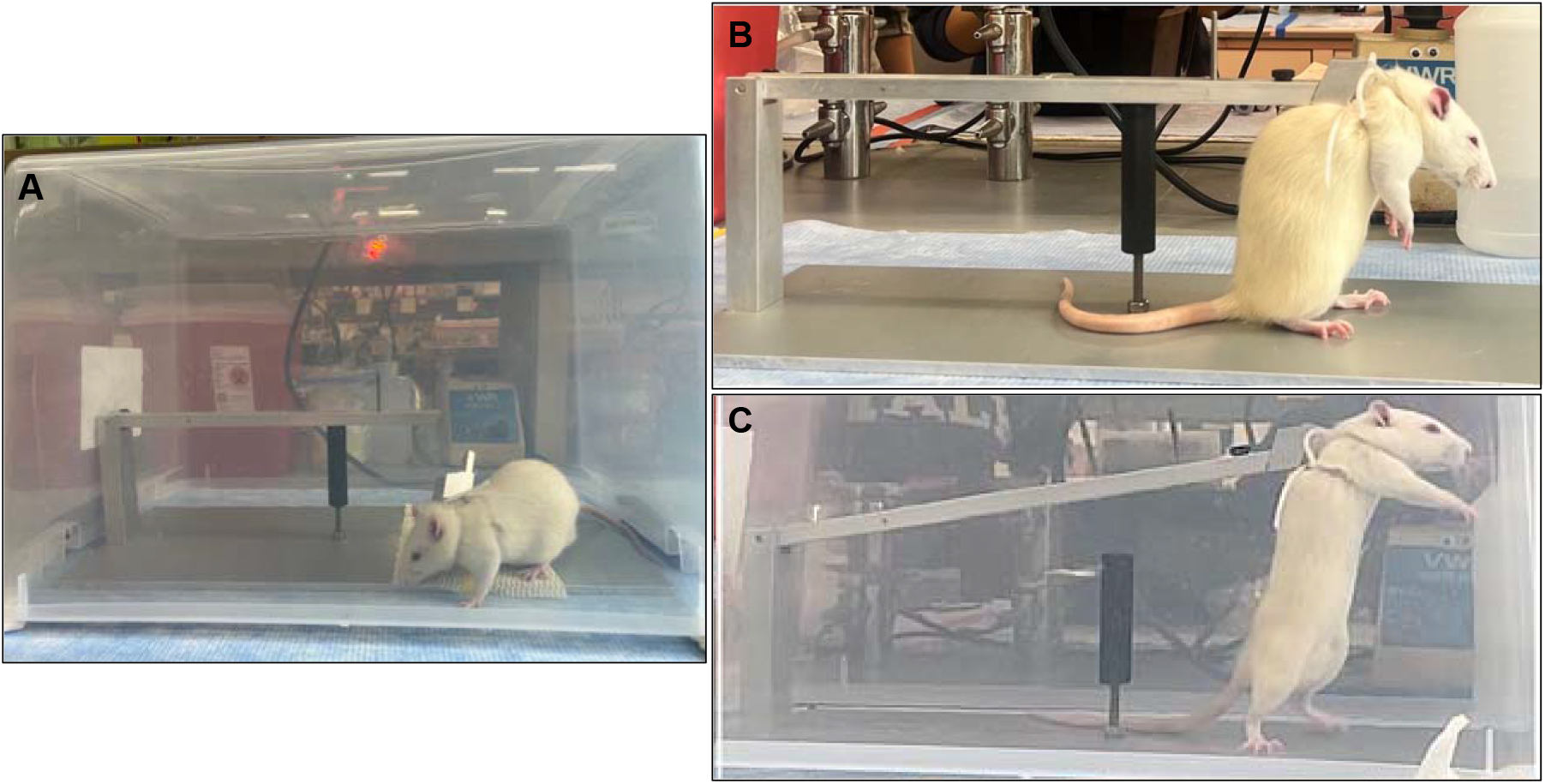
Generation of a resistance exercise model in rats. (A) Representative photo of day 3-4 of during the 7–10-day exercise acclimation period. The flexible harness is strapped to the rat. The rat is left to freely roam the squat apparatus in a confined space. (B) The rat secured onto the squat apparatus in the ‘rest’ position (full knee flexion). (C) Rat harnessed in the squat apparatus in a full knee extension position.

## DISCUSSION

BFR is a promising tool used to increase muscle mass and attenuate muscle atrophy in individuals experiencing muscle loss, including patients recovering from orthopedic surgery. To develop appropriate guidelines for the use of BFR-exercise in a rehabilitation setting, a comprehensive understanding of its underlying physiologic mechanisms is crucial. This necessitates the use of a suitable animal model. In this methods paper, we describe a comprehensive protocol establishing a rat model of BFR-exercise in a post-surgical rehabilitative setting. The protocol includes ACL reconstruction (ACL-R) surgery to establish an atrophic muscle state, occlusive cuff placement to induce transient intramuscular hypoxia, loaded knee extension exercise that isolates hindlimb musculature, and *in vivo* muscle biopsy methods for subsequent tissue analysis.

Here, we created a protocol to surgically transect and reconstruct the ACLs of rat hindlimbs bilaterally in a single procedure using a commercially available copolymer graft. The use of engineered co-polymer grafts to re-establish ligament function in rat ACL-R surgery has been previously described with varying efficacy^42,44,55–59^. Most commonly, these grafts are constructed in a laboratory with the infrastructure and skillsets to build and test these materials. This environment is often limited or inaccessible. FlexBand is commercially available and approved for use in humans to augment soft tissue reconstruction procedures for foot and ankle reconstruction and sports medicine applications^60,61^. Previous research has tested a similar co-polymer product in large animals^62,63^. To our knowledge, the FlexBand product has not been used as an ACL replacement in rats. Here, we show successful integration and recapitulation of the functional properties of the ACL in the operative limbs using the FlexBand. We also show ∼26% reduction in quadriceps weight two-weeks following ACL-R in rats, demonstrating that ACL-R is a viable model of muscle loss in rats. Considering that sarcopenia is an independent predictor of a 37% increase in mortality risk across studies in elderly people and individuals with cancer, cardiovascular, liver, lung, and renal diseases^64^, there is an urgent need for new models that recapitulate acute muscle loss and which can be used to test methods to reverse it.

Although ACL tears, and subsequent reconstructive surgery, most commonly occur on a single leg in humans, we chose to perform bilateral ACL transection and reconstruction to best integrate into studies of BFR-exercise in awake rats. Bilateral ACL reconstruction allows for access to a larger amount of experimental tissue and avoids potentially confounding variables from the non-surgical contralateral leg during metabolic or physiologic analyses. Further, it avoids compensation by the non-surgical limb, as it is difficult to isolate a single hindlimb through physical manipulation during strength exercise in an awake rat.

To determine the occlusion pressures necessary to distinguish therapeutic versus non-therapeutic BFR in rats, we used the Visualsonics Vevo 3100 doppler ultrasound paired with the UHF46x transducer. We measured blood flow (mm/s) and vessel diameter (mm) over the external iliac artery while gradually increasing external occlusion to the following pressures using a 1.6cm digit cuff (Hokason, Inc): 0, 40, 60, 80, 100, and 120 mmHg. We observed a 24% vessel occlusion at 80 mmHg, which is below the therapeutic range (>40% occlusion). Our findings are not entirely consistent with other studies that use external cuffs to interrogate vascular changes. Yoshikawa et al. used the same external occlusion cuff as presented in this paper to evaluate the effects of BFR on muscle growth in neuromuscular electro-stimulated planta pedis and soleus muscles in anesthetized rats. The group observed a 22% occlusion of blood flow to the planta pedis muscle and a 54% vessel occlusion of blood flow to the soleus muscle at 80 mmHg and no further occlusion with increasing external pressure^47^. Tanaka et al. observed a 62% occlusion in blood flow to the planta pedis muscle and 22% occlusion in blood flow to the soleus muscle at 80 mmHg using the same cuffs^65^. This discrepancy may exist because of methodological differences: the aforementioned studies measure blood flow into muscle groups distal to the occlusion cuff (planta pedis and soleus), however, it is not specified which vessels were probed in these experiments.

In our experiments, we use a high-sensitivity ultrasound to visualize and directly measure hindlimb vasculature in rats, compared to existing studies that probe muscle groups to determine blood flow. When attempting to probe the femoral artery or any distal vessel, we found it difficult to achieve accurate probe and cuff placement simultaneously as the cuff covered most of the surface area of the rat’s hindlimb above the knee. Further, any attempt to measure vasculature distal to the knee joint was limited due to small vessel size. We chose to probe over the external iliac artery, which we identified above the occlusion cuff. Probing of this more proximal vessel allowed for better views and more accurate reading of blood flow and vessel diameter. We may have observed sub-therapeutic pressures for a few reasons. First, the location on the external iliac artery where measurements were collected is just distal to the bifurcation point of the external and iliac artery. It is possible that the readings we measured were affected by turbulent blood flow. Second, depending on the time it takes to orient the animal and identify the vessel, vascular changes, including dilation of the artery, may occur in settings of prolonged exposure to anesthesia. Given the challenges expressed here and the limitations of existing studies, accurate determination of thresholds for BFR will be necessary to provide a foundation for future experiments utilizing the external cuff.

Use of a digit cuff (1.6 and 1.9 cm) has been previously described to establish BFR-exercise via electro-stimulated muscle contractions in anesthetized rats^47^. This methodology bypasses the challenges of placing the cuff onto an awake animal in preparation for BFR-exercise, like cuff material and animal positioning. We found that the 1.6cm cuff size is appropriate for the upper hindlimb of adult rats as small as ∼300-400g. While attempting to adapt the 1.6cm cuff to our model of BFR-exercise in the awake rats, we found it difficult to secure and stabilize the cuff during *in vivo* experiments. We found that applying the cuff lined with a thin strip of double-sided tape after removing the fur from the hindlimb worked best to maintain cuff position and function during strength exercise. We attached the cuff after the rat was strapped onto the exercise apparatus and under gentle restraint by hand while in the machine. This allows for targeted induction and maintenance of BFR in an awake, exercising rat.

As mentioned above, it is difficult to isolate the limbs involved in the BFR intervention during exercise, as in modalities like tail-weighted ladder climbing and incline treadmill walking. Although technically advantageous, these methods do not directly replicate the clinical use of BFR-exercise in rehabilitation settings. In this paper, we described use of a squat apparatus (adapted from Tamaki et al.), which allows for isolated full knee flexion and extension of the surgical hindlimbs. Other groups have also adapted this apparatus to model resistance exercise in rats^66–68^. However, there has not been a direct application of this resistance exercise model in combination with BFR in rats. We configured flexible harnesses to the machine both for optimal rodent comfort and the ability to access vascular lines for experimental infusions or blood collection following BFR-exercise. Other studies using this apparatus also detail a training protocol that includes distinct sets and repetitions (*eg*. 4 sets of 12 repetitions)^66^. However, we found it difficult to reproducibly enforce these training protocols. To mitigate this, we extended our acclimation period as needed for each animal to reduce sporadic movements. Second, we prompted knee extension with compressed air targeted to the rectum as opposed to tail shocks to encourage full range of motion movements and reduce large stress responses. Third, we counted repetitions based on the quality of movement (*eg*. full knee extension and return to full knee flexion as one repetition). This methods description allows for variability in the number of sets and repetitions and/or the duration of exercise performed by the exercising animal. The nature of the apparatus also allows for modification of squat depth and loaded weight. Full repetitions, whether self-motivated or encouraged with compressed air, can be manipulated by modifying of squat depth and loaded weight.

There are several strengths of the current study. This methods paper offers: (1) a clinically relevant protocol in an accessible animal model, (2) an option to address the limitations of previous studies that use non-physiologic models of resistance exercise in anesthetized rats, where we describe a method to perform strength training exercise awake rats, and (3) a comprehensive protocol targeted for use in mechanistic and physiologic studies of BFR-exercise.

Despite these strengths, there are a few weaknesses. The experiments included in this paper include relatively small sample sizes. Further, only male rats were used in these experiments, although, in humans, females are at higher risk for ACL tears^69^. In our attempt to generate therapeutic BFR, we observed ∼24% occlusion of the external iliac artery. As the lower threshold of therapeutic occlusion pressure is described to be 40%^70^, we have not yet documented major occlusion. Given that our observations are from a single rat, repeated measurements are warranted. Further, blood flow occlusion is measure above, not below, the occlusion cuff. This may not accurately reflect the downstream vascular effects of the targeted ischemia produced by partial blood flow occlusion. Further, the set-up of our BFR and exercise model, especially inducing squats with compressed air, is likely to induce a stress response in the rats, which can affect tolerability of the intervention and confound any metabolic and behavioral outputs of these tests. Lastly, the bilateral ACL-R model that we describe requires use of a synthetic allograft of the ACL, which does not exactly mimic an autograft of the ACL.

In conclusion, we demonstrate a comprehensive, clinically relevant protocol to establish BFR-exercise in an awake rat model. This protocol includes generating a muscle loss state via bilateral ACL-R, determining targeted blood flow occlusion pressures, and performing weighted hind-limb knee extension exercises with BFR. These methods can be used for further application in mechanistic and physiologic studies of BFR, resistance exercise, or combined BFR-exercise.

## ACKNOWLEDGMENTS

The authors would like to thank the Yale Machine Shop for their technical contributions to creation of the squat apparatus described in this manuscript. The authors would also like to thank Artelon (Sandy Springs, GA) for their generosity in supplying the FlexBand product for use in these experiments. This publication was made possible by CTSA Grant Number UL1 TR001863 from the National Center for Advancing Translational Science (NCATS), a component of the National Institutes of Health (NIH). Its contents are solely the responsibility of the authors and do not necessarily represent the official views of NIH.

## CONFLICTS OF INTEREST

None.

## Notes

### Competing Interest Statement

The authors have declared no competing interest.

